# A Connectome of the Male *Drosophila* Ventral Nerve Cord

**DOI:** 10.1101/2023.06.05.543757

**Authors:** Shin-ya Takemura, Kenneth J Hayworth, Gary B Huang, Michal Januszewski, Zhiyuan Lu, Elizabeth C Marin, Stephan Preibisch, C Shan Xu, John Bogovic, Andrew S Champion, Han SJ Cheong, Marta Costa, Katharina Eichler, William Katz, Christopher Knecht, Feng Li, Billy J Morris, Christopher Ordish, Patricia K Rivlin, Philipp Schlegel, Kazunori Shinomiya, Tomke Stürner, Ting Zhao, Griffin Badalamente, Dennis Bailey, Paul Brooks, Brandon S Canino, Jody Clements, Michael Cook, Octave Duclos, Christopher R Dunne, Kelli Fairbanks, Siqi Fang, Samantha Finley-May, Audrey Francis, Reed George, Marina Gkantia, Kyle Harrington, Gary Patrick Hopkins, Joseph Hsu, Philip M Hubbard, Alexandre Javier, Dagmar Kainmueller, Wyatt Korff, Julie Kovalyak, Dominik Krzemiński, Shirley A Lauchie, Alanna Lohff, Charli Maldonado, Emily A Manley, Caroline Mooney, Erika Neace, Matthew Nichols, Omotara Ogundeyi, Nneoma Okeoma, Tyler Paterson, Elliott Phillips, Emily M Phillips, Caitlin Ribeiro, Sean M Ryan, Jon Thomson Rymer, Anne K Scott, Ashley L Scott, David Shepherd, Aya Shinomiya, Claire Smith, Natalie Smith, Alia Suleiman, Satoko Takemura, Iris Talebi, Imaan FM Tamimi, Eric T Trautman, Lowell Umayam, John J Walsh, Tansy Yang, Gerald M Rubin, Louis K Scheffer, Jan Funke, Stephan Saalfeld, Harald F Hess, Stephen M Plaza, Gwyneth M Card, Gregory SXE Jefferis, Stuart Berg

## Abstract

Animal behavior is principally expressed through neural control of muscles. Therefore understanding how the brain controls behavior requires mapping neuronal circuits all the way to motor neurons. We have previously established technology to collect large-volume electron microscopy data sets of neural tissue and fully reconstruct the morphology of the neurons and their chemical synaptic connections throughout the volume. Using these tools we generated a dense wiring diagram, or connectome, for a large portion of the *Drosophila* central brain. However, in most animals, including the fly, the majority of motor neurons are located outside the brain in a neural center closer to the body, i.e. the mammalian spinal cord or insect ventral nerve cord (VNC). In this paper, we extend our effort to map full neural circuits for behavior by generating a connectome of the VNC of a male fly.

## 1 Introduction and Motivation

Connecting structure to function is the ultimate goal when seeking a mechanistic understanding of a biological system. The fruit fly, *Drosophila melanogaster*, has emerged as a powerful model system for studying how the nervous system generates a wide range of complex motor and cognitive behaviors such as courtship, navigation, associative memory, foraging for food and escaping from predators. The compact size of the fly nervous system, together with genetic, physiological, and other experimental tools make possible the characterization, manipulation, and functional assessment of individual neurons and synapses throughout the central nervous system (CNS). In particular, one class of tools that have greatly accelerated the functional dissection of neural circuits in the fly is connectomes, complete wiring diagrams and connectivity matrices generated from large-volume electron microscopy datasets of particular brain regions ([Zheng et al., 2018]; reviewed in [Galili et al., 2022]; [Scheffer et al., 2020]; [Phelps et al., 2021]). Currently, densely-reconstructed (with every neuron traced and proofread) connectomes exist for several columns in the optic lobe and a large portion of a female fly central brain (hemibrain; [Scheffer et al., 2020]) An effort to reconstruct the entire female fly brain is nearing completion [https://flywire.ai/].

The resolution of electron microscopy enables the reconstruction of both the detailed structure of neurons as well as identification of synapses, from which connectivity and strength can be resolved, allowing for rich functional hypotheses generation and testing. The power of connectomes to provide mechanistic insights into critical central brain computations has already been demonstrated in processes as diverse as calculating the direction of visual motion (see for example [Shinomiya et al., 2022]) or maintaining an internal representation of heading (see for example, [Hulse et al., 2021]), and implementing reinforcement rules for associative learning (see for example [Li et al., 2020]). However, it remains unclear how these critical computations actually guide actions of the fly. This is because existing densely-reconstructed connectomes do not provide information about premotor circuits or the motor neurons themselves, which are the elements of the nervous system that extend to the periphery and activate muscles that move the limbs about their various joints. As in mammals, in the fly these motor neurons are mostly contained within a portion of the nervous system that is separate from the brain and located in the animal’s body: the ventral nerve cord (VNC, analogous to the mammalian spinal cord). A partially-reconstructed VNC in the female fly [Phelps et al., 2021] has already demonstrated how important elucidating the structure and connectivity of the pre-motor circuits in the nerve cord is for understanding how commands from the brain are translated into behavior.

Here we present the first densely-reconstructed connectome for the VNC of a fly, which we refer to as MANC, the Male Adult Nerve Cord. The position of the VNC in the fly is shown in Fig. 1. The VNC receives input from the brain in the form of descending neurons, a population of neurons that enter as a group through the cervical connective, a narrow bundle of parallel fibers that links the ventral area of the brain to the anterior aspect of the VNC. The VNC is known to have a layered organization, with motor neurons for wing muscles and leg muscles occupying the dorsal and ventral layers respectively. In addition to these descending motor commands, the VNC also receives sensory input directly from a wide range of body-centric sensory systems and has complex local circuits that integrate the descending and sensory inputs to influence outgoing commands to the motor neurons. We chose to reconstruct the VNC of a male fly as there are sex-specific motor programs displayed only by males that are critical for courtship and mating, including song production and copulation behaviors. This dataset also complements a partially proofread female VNC connectome ([Phelps et al., 2021]; [Azevedo et al., 2022]), allowing for future detailed comparison of male and female premotor circuits.

**Figure 1:**
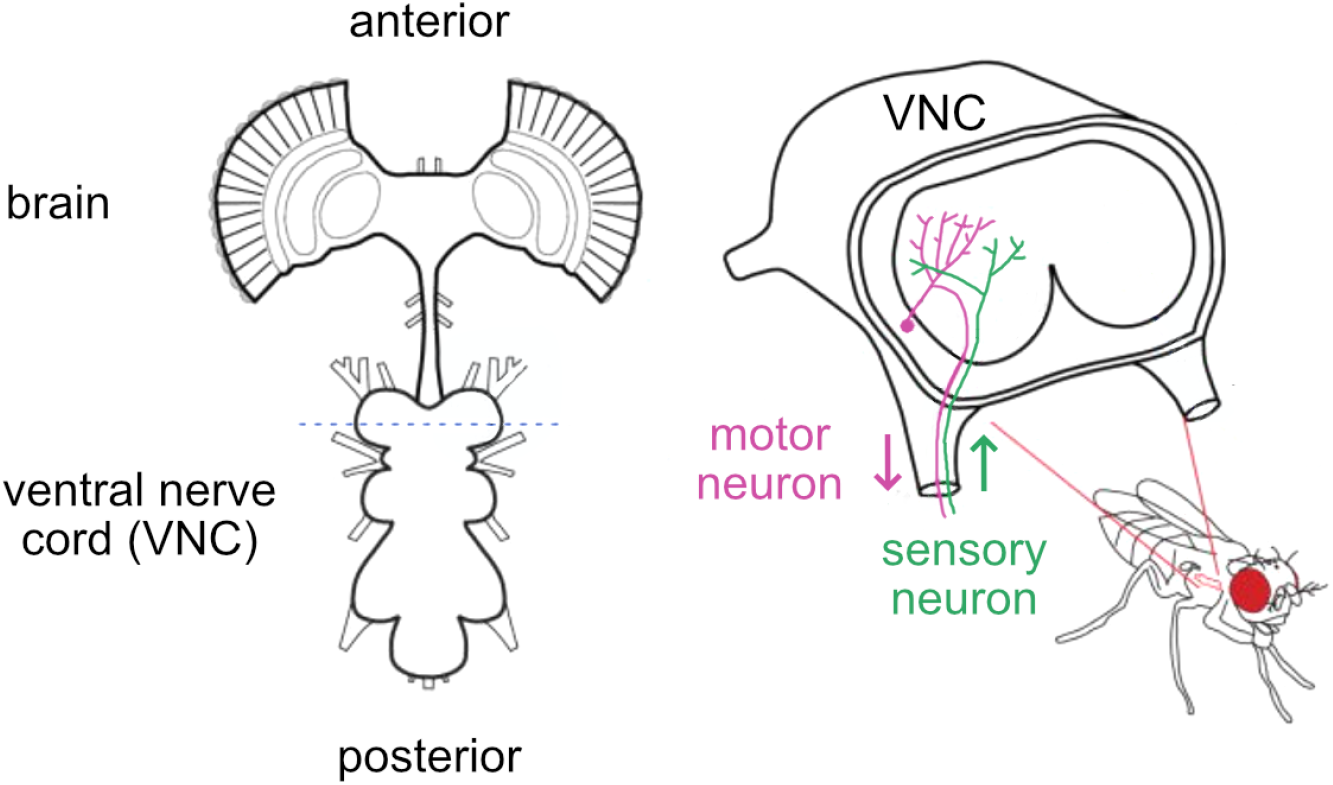
The ventral nerve cord, and its location within the fly’s body. Source: John.Tuthill, https://commons.wikimedia.org/w/index.php?curid=90902274

**Figure 2:**
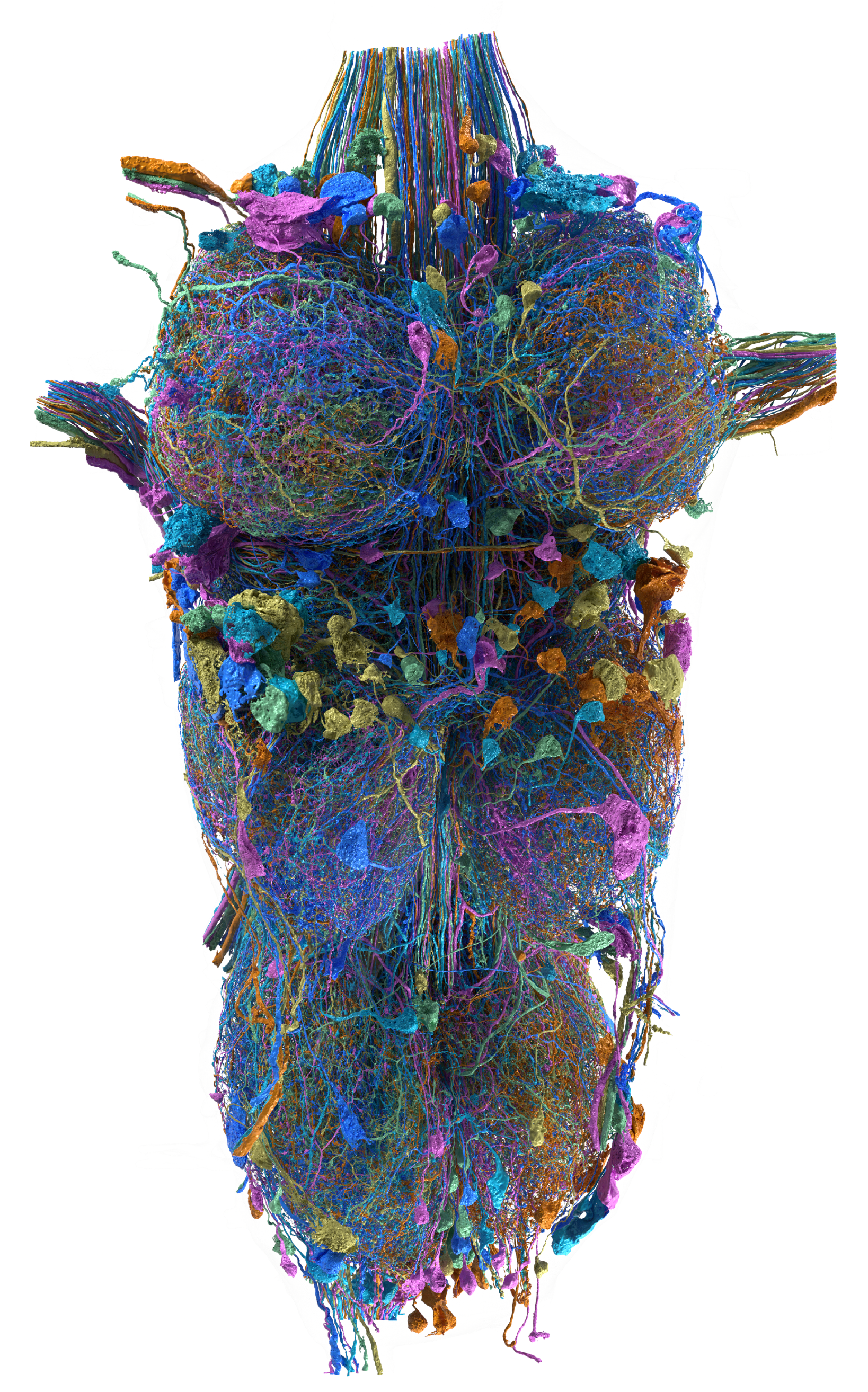
A selection of reconstructed neurons. The neurons shown here comprise 4% of all reconstructed neurons in the MANC. (Ventral view.)

Our VNC sample contains roughly 23 thousand traced neurons, 10 million TBars, 74 million PSDs, and 44m of neuronal cable. We divide the VNC into a set of regions of interest (ROIs) based on established anatomical references (Fig. 3, [Court et al., 2020]). These ROIs comprise two kinds, neuropils and nerves. The reconstructed neuropils are shown in Table 1 and the nerves in Table 2. The nerves have many fewer synapses and are much less well reconstructed, largely due to damage from truncation during sample preparation.

**Table 1:** Neuropils contained and defined in the ventral nerve cord, following the naming conventions of [Court et al., 2020] with the addition of (R) and (L) to specify the side of the soma for that region. Counts are the number of objects found within the region; percentage is the fraction in traced neurons. Connection completion percentage is the fraction of all TBar-PSD pairs where both are in traced neurons. LegNP = leg neuropil; ANm = abdominal neuromeres; mVAC=medial ventral association center; Ov = ovoid neuropil; NTct = neck tectulum; WTct=wing tectulum; HTct = haltere tectulum; IntTct = intermediate tectulum; LTct = lower tectulum

| ROI | TBars |  | PSDs |  | Connection |
| --- | --- | --- | --- | --- | --- |
|  | count | % | count | % | comp. % |
| LegNp(T1)(L) | 1132044 | 86 | 8324620 | 39 | 35 |
| LegNp(T1)(R) | 1128824 | 87 | 8151463 | 42 | 37 |
| LegNp(T2)(L) | 1014038 | 87 | 7518749 | 43 | 39 |
| LegNp(T2)(R) | 1031882 | 89 | 7798126 | 48 | 43 |
| LegNp(T3)(L) | 1154981 | 86 | 8917873 | 38 | 33 |
| LegNp(T3)(R) | 1167221 | 90 | 8747460 | 45 | 40 |
| ANm | 945949 | 96 | 6909412 | 54 | 52 |
| mVAC(T1)(L) | 33737 | 87 | 205525 | 51 | 44 |
| mVAC(T1)(R) | 38512 | 90 | 234131 | 56 | 51 |
| mVAC(T2)(L) | 22528 | 80 | 147423 | 54 | 44 |
| mVAC(T2)(R) | 24219 | 90 | 158243 | 62 | 56 |
| mVAC(T3)(L) | 34373 | 87 | 225916 | 58 | 52 |
| mVAC(T3)(R) | 35765 | 90 | 232019 | 60 | 55 |
| Ov(L) | 193036 | 98 | 1352587 | 62 | 60 |
| Ov(R) | 202970 | 97 | 1349857 | 63 | 62 |
| NTct(UTct-T1)(L) | 110522 | 97 | 755793 | 36 | 35 |
| NTct(UTct-T1)(R) | 122861 | 96 | 801679 | 36 | 35 |
| WTct(UTct-T2)(L) | 290650 | 97 | 1753286 | 55 | 54 |
| WTct(UTct-T2)(R) | 286207 | 97 | 1670457 | 55 | 53 |
| HTct(UTct-T3)(L) | 248578 | 97 | 1731843 | 47 | 46 |
| HTct(UTct-T3)(R) | 258975 | 97 | 1763739 | 47 | 46 |
| IntTct | 408419 | 96 | 2844569 | 45 | 43 |
| LTct | 359394 | 95 | 2414756 | 51 | 49 |

**Table 2:**
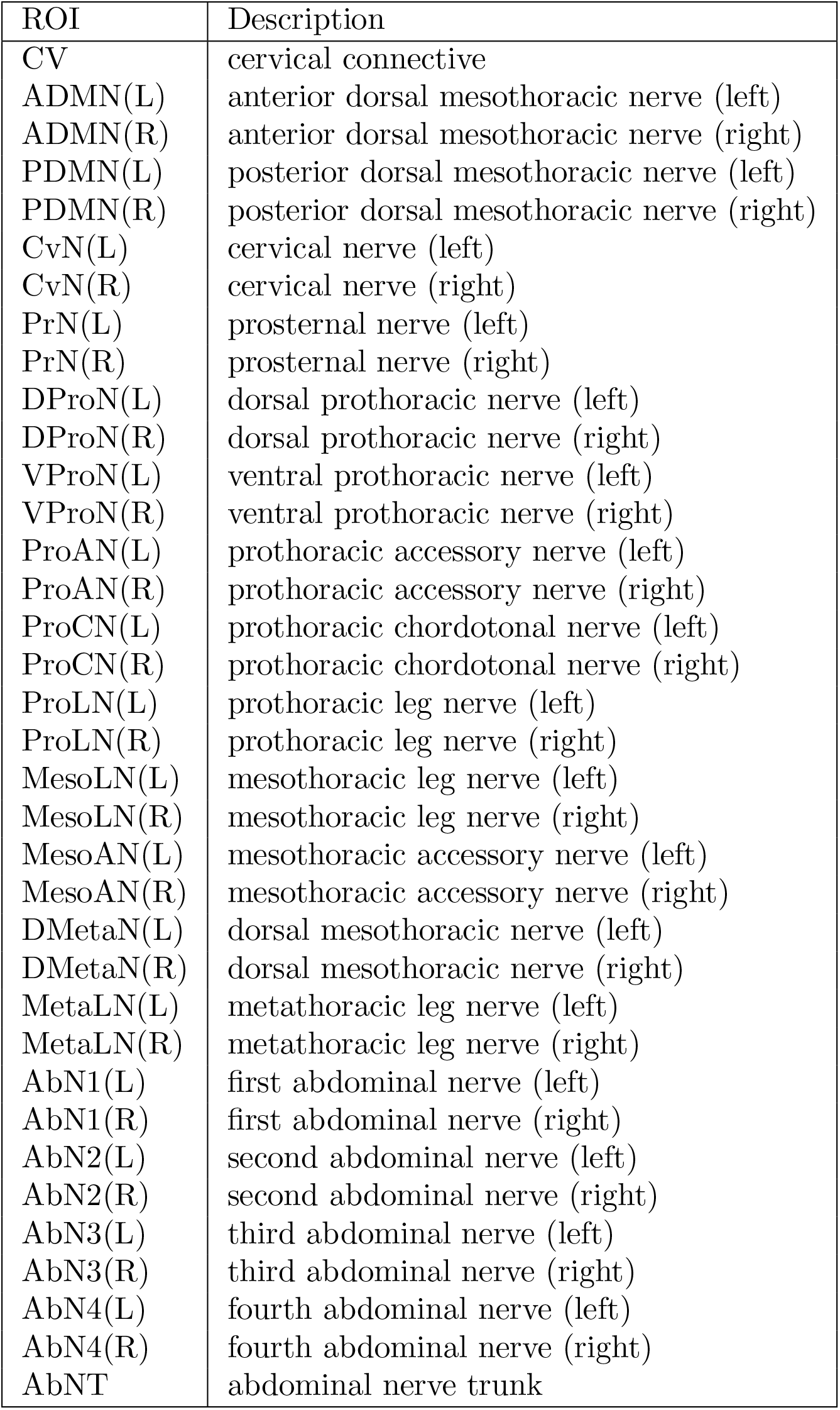
Nerve regions contained and defined in the ventral nerve cord, following the naming conventions of [Court et al., 2020] with the addition of (R) and (L) to specify the side of the soma for that region.

| ROI | Description |
| --- | --- |
| CV | cervical connective |
| ADMN(L) | anterior dorsal mesothoracic nerve (left) |
| ADMN(R) | anterior dorsal mesothoracic nerve (right) |
| PDMN(L) | posterior dorsal mesothoracic nerve (left) |
| PDMN(R) | posterior dorsal mesothoracic nerve (right) |
| CvN(L) | cervical nerve (left) |
| CvN(R) | cervical nerve (right) |
| PrN(L) | prosternal nerve (left) |
| PrN(R) | prosternal nerve (right) |
| DProN(L) | dorsal prothoracic nerve (left) |
| DProN(R) | dorsal prothoracic nerve (right) |
| VProN(L) | ventral prothoracic nerve (left) |
| VProN(R) | ventral prothoracic nerve (right) |
| ProAN(L) | prothoracic accessory nerve (left) |
| ProAN(R) | prothoracic accessory nerve (right) |
| ProCN(L) | prothoracic chordotonal nerve (left) |
| ProCN(R) | prothoracic chordotonal nerve (right) |
| ProLN(L) | prothoracic leg nerve (left) |
| ProLN(R) | prothoracic leg nerve (right) |
| MesoLN(L) | mesothoracic leg nerve (left) |
| MesoLN(R) | mesothoracic leg nerve (right) |
| MesoAN(L) | mesothoracic accessory nerve (left) |
| MesoAN(R) | mesothoracic accessory nerve (right) |
| DMetaN(L) | dorsal mesothoracic nerve (left) |
| DMetaN(R) | dorsal mesothoracic nerve (right) |
| MetaLN(L) | metathoracic leg nerve (left) |
| MetaLN(R) | metathoracic leg nerve (right) |
| AbN1(L) | first abdominal nerve (left) |
| AbN1(R) | first abdominal nerve (right) |
| AbN2(L) | second abdominal nerve (left) |
| AbN2(R) | second abdominal nerve (right) |
| AbN3(L) | third abdominal nerve (left) |
| AbN3(R) | third abdominal nerve (right) |
| AbN4(L) | fourth abdominal nerve (left) |
| AbN4(R) | fourth abdominal nerve (right) |
| AbNT | abdominal nerve trunk |

**Figure 3:**
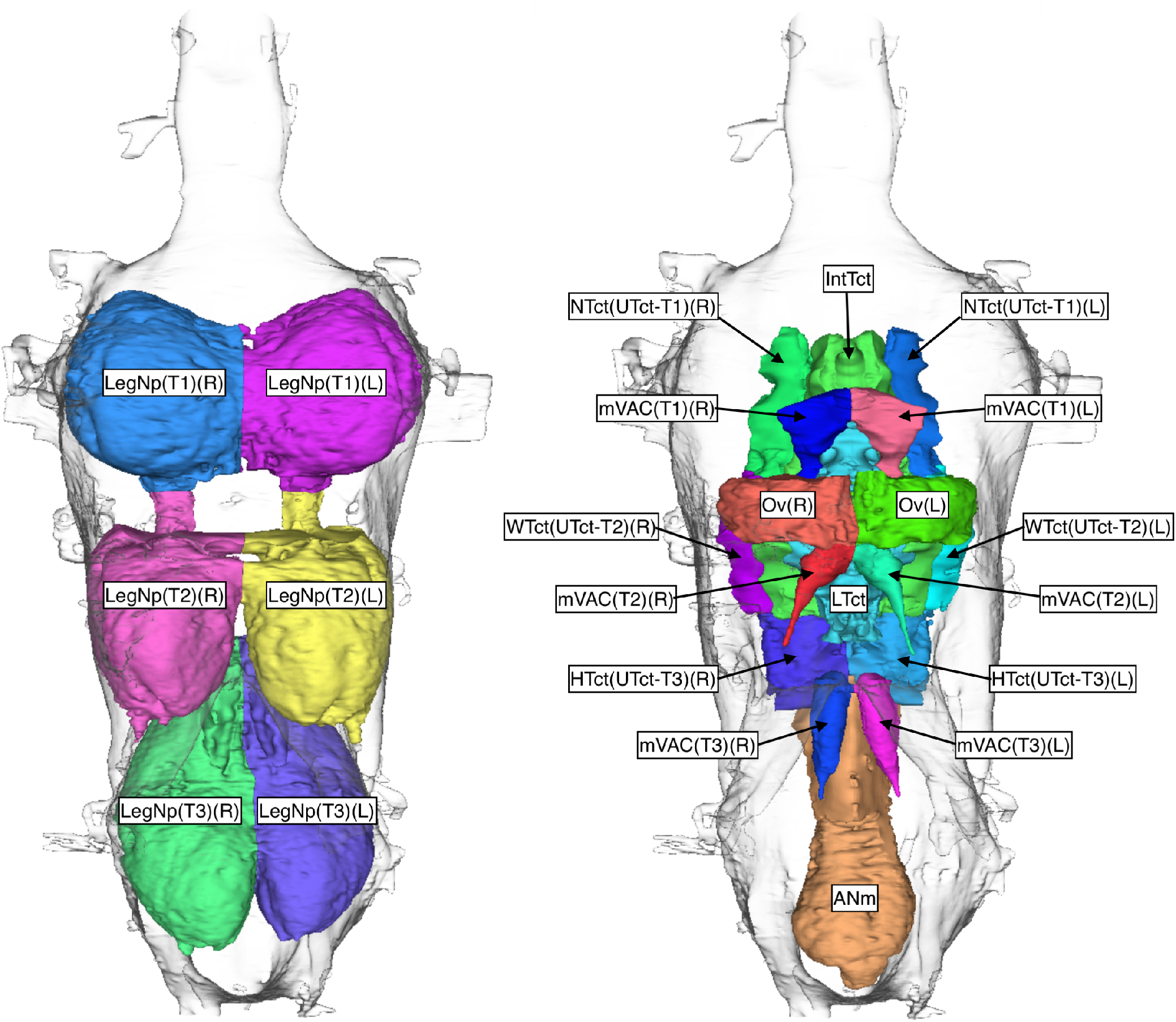
Neuropil compartments in the ventral nerve cord. (Ventral views.)

This paper describes the methods used to obtain the data set. Two companion papers provide additional biological information: [Marin et al., 2023] describe the annotation of neurons across the VNC and their grouping into cell types; and (2) Cheong, Eichler, Stürner et al. describe the organization of descending- to-motor pathways including the identification of descending and motor neurons. An overview of introductory materials, exported datasets, and introductory material for the MANC connectome can be found at https://www.janelia.org/manc-connectome.

The VNC connectome is publicly available to anyone with a google account at https://neuprint.janelia.org, dataset MANC. This web address brings up the application ‘NeuPrint’, which allows many different queries about the VNC data. It is intended to be largely self-explanatory for those already familiar with *Drosophila* anatomy. A tutorial video is available at https://www.youtube.com/watch?v=FlpMlS9lixk. In addition, there are Application Program Interfaces (APIs) that offer access to the connectome from the computer languages *Python, R*, and *Cypher*. This includes a malevnc R package specialized for this dataset which was used for the majority of analyses in the two companion papers (https://natverse.org/malevnc; see [Marin et al., 2023] for further details). The neuPrint database includes the full set of annotations for intrinsic and sensory neurons described by [Marin et al., 2023] as well as descending and motor neurons identified by Cheong, Eichler, Stürner et al. Researchers making use of this annotation information should cite the respective source study.

## 2 Methods

### 2.1 Sample preparation

In general, the sample preparation methods were similar to those used in [Scheffer et al., 2020] and described in [Lu et al., 2019]. Five day old males from a cross between wild-type Canton S strain G1 x w^1118^ were raised on a 12 hour day/night cycle and dissected 1.5 hours after lights-on. The main difference from previous work occurred during this dissection step. Here we dissected the entire central nervous system as a unit, comprising the brain, the ventral nerve cord, and the neck. In this work, however, only the ventral nerve cord was imaged and reconstructed.

Samples were fixed in aldehyde fixative, enclosed in bovine serum albumin (BSA) and post-fixed in osmium tetroxide. Samples were then treated with potassium ferricyanide and incubated with aqueous uranyl acetate followed by lead aspartate staining. A Progressive Lowering Temperature (PLT) dehydration procedure was applied to the samples. After PLT and low temperature incubation with an ethanol based UA stain, the samples were infiltrated and embedded in Epon (Poly/Bed 812;Luft formulations). Collectively these methods optimize morphological preservation, allow full-brain preparation without distortion, and provide increased staining intensity which in turn enables faster FIB-SEM imaging. Each completed sample was examined by x-ray CT imaging (Zeiss Versa 520) to check for dissection damage and other potential flaws, and assess the quality of the staining.

### 2.2 Hot knife cutting

The VNC sample is *>*500 µm long and, in places, as large as 250 µm diameter, making it too large to be practically FIB-SEM imaged without introducing milling artifacts. Using the approach developed during the hemibrain project [Scheffer et al., 2020] of dividing the sample into thick slices prior to imaging, we subdivided the VNC lengthwise into 25 micron thick slabs; this also allowed imaging individual slabs in parallel across multiple FIB-SEM machines. The heavy-metal stained VNC (surrounded by BSA) was embedded in a resin block such that its long axis was perpendicular to the blockface. This block was imaged by x-ray CT (Zeiss Versa 520) and trimmed into a fin shape whose width was minimized (600 microns) to reduce forces on the diamond knife during sectioning. We used our previously described ‘hot-knife’ ultrathick sectioning procedure [Hayworth et al., 2015] which uses a heated, oil-lubricated diamond knife (Diatome 25^*◦*^ angle Cryo-Dry diamond knife, 90 ^*◦*^C knife set point temperature, 0.1 mm/s cutting speed) to create a total of 26 slabs, 25 of which were imaged and used in the final reconstruction. Each oil-covered slab was imaged by light microscopy for quality control, and then flat embedded in Durcupan resin against a 25 micron thick PET backing film. Individual slabs were then trimmed out of the flat embedding resin using a scalpel, glued onto metal tabs suitable for mounting in the FIB-SEM, and laser trimmed to remove most non-tissue regions surrounding the VNC tissue to allow for efficient use of FIB milling time. Each mounted slab was then x-ray CT imaged at 0.67 micron voxel size to check sample quality and to establish a scale factor for Z-axis cutting by FIB. The resulting slabs were FIB-SEM imaged separately and the resulting volume datasets were stitched together computationally as discussed in the section on alignment.

### 2.3 Imaging

The twenty-five VNC slabs were imaged using seven customized FIB-SEM systems in parallel over a period of six months. Unlike the FIB-SEM machines used for the hemibrain project [Scheffer et al., 2020], this new platform replaced the FEI Magnum FIB column with the Zeiss Capella FIB column to improve the precision and stability of FIB milling control [Xu et al., 2021]. Specifically, FIB milling was carried out by a 15-nA 30 kV Ga ion beam with an 8-nm step size. SEM images were acquired at 8-nm XY pixel size at 3 MHz using a 3-nA beam with 1.2 kV landing energy. Specimens were grounded at 0V to enable both secondary and backscattered electrons collection.

### 2.4 Alignment

Alignment followed the methods of hemibrain[Scheffer et al., 2020] but with substantial improvements. A pipeline based upon the ‘*render*’ web services, together with special code for image alignment across hot knife seams, was used to align and reconstruct the FIB-SEM images into 3D volumes for each of the 25 slabs (sections 2 to 26 of the original sample). In addition to SIFT[Lowe, 2004], geometric local descriptor matching[Preibisch et al., 2010] was employed for feature matching in resin-heavy areas, was incorporated to automatically solve problematic areas where imaging was interrupted. As with the hemibrain, milling thickness variations in the aligned series were compensated using a modified version of the method described by [Hanslovsky et al., 2017].

To straighten the aligned hot-knife slabs prior to final stitching, the hemibrain methods for surface finding adapted from Kainmueller[Kainmueller et al., 2008] were further improved including an ImgLib2-based[Pietzsch et al., 2012] distributed cost and graph-cut computation as well as the development of a Big-DataViewer[Pietzsch et al., 2015] based tool for interactively fixing remaining issues. As with the hemibrain, the series of flattened slabs was then stitched using a custom, distributed method for large-scale deformable registration to account for deformations introduced during hot-knife sectioning. These volumes were then contrast adjusted using local contrast normalization (CLLCN) prior to segmentation.

These techniques are described in more detail in (Preibisch, Trautman and Saalfeld, 2023, in preparation). The code used for the ‘*render*’ web services can be found at https://github.com/saalfeldlab/render and the code used for surface finding and hot-knife image stitching is available at https://github.com/saalfeldlab/hot-knife.

### 2.5 Segmentation

To reconstruct neurites from the EM imagery, we used multi-scale flood-filling networks[Januszewski et al., 2018] as in the hemibrain paper. However, the use of lower resolution models (operating on 32×32×32 nm^3^ and 16×16×16 nm^3^ voxels) trained on 111 manually proofread neurites from the hemibrain allowed us to segment the CLAHE-normalized images of the VNC directly without CycleGAN normalization in the interior of hotknife tabs. Instead, at 8×8×8 nm^3^ and 16×16×16 mm^3^ resolution only the image content within 5 voxels around every hot knife tab interface was replaced with that generated by segmentation-enhanced cycle-consistent generative adversarial networks (SECGANs) [Januszewski and Jain, 2019] to improve neurite contiguity. We did not perform such replacement in the 32×32×32 nm^3^ imagery as we could not observe a measurable improvement from this procedure.

Compared to the hemibrain, we applied some additional constraints during agglomeration. We manually generated point annotations for cell bodies, neck fibers, and nerve bundles, which were used to forbid merges between the corresponding segments. We also applied heuristics that disallow merges between any two objects that have accreted voxels beyond a certain threshold size (10M voxels) during agglomeration with the 8mn model. These constraints help prevent the creation of very large bodies containing several cells merged together. Such cells are very difficult and time consuming to split manually, and preventing their formation paid large dividends in proofreading time.

Similarly to the hemibrain, we also used a separate tissue classification model with a 5-class label space (out of bounds, trachea, glia, cell bodies, neuropil). Voxels classified as out of bounds, trachea or glia were masked during FFN segmentation. Segments with a dominant ”neuropil” voxel classification were considered for agglomeration with 32 nm, 16 nm and 8 nm models, whereas segments with a dominant ”soma” classification were only considered for agglomeration with 32 nm and 16 nm models.

### 2.6 Synapse identification

Synapse prediction was performed as described in hemibrain project[Scheffer et al., 2020]. Through careful selection of training and validation data spanning the VNC ROIs, we were able to achieve desired performance with a single network for detecting pre-synaptic T-bars, and did not require the cascaded approach as used in previous work. The post-synaptic partners were found using the procedures used for the hemibrain[Scheffer et al., 2020]. The precision-recall plots for synapse identification are shown in Fig. 4.

**Figure 4:**
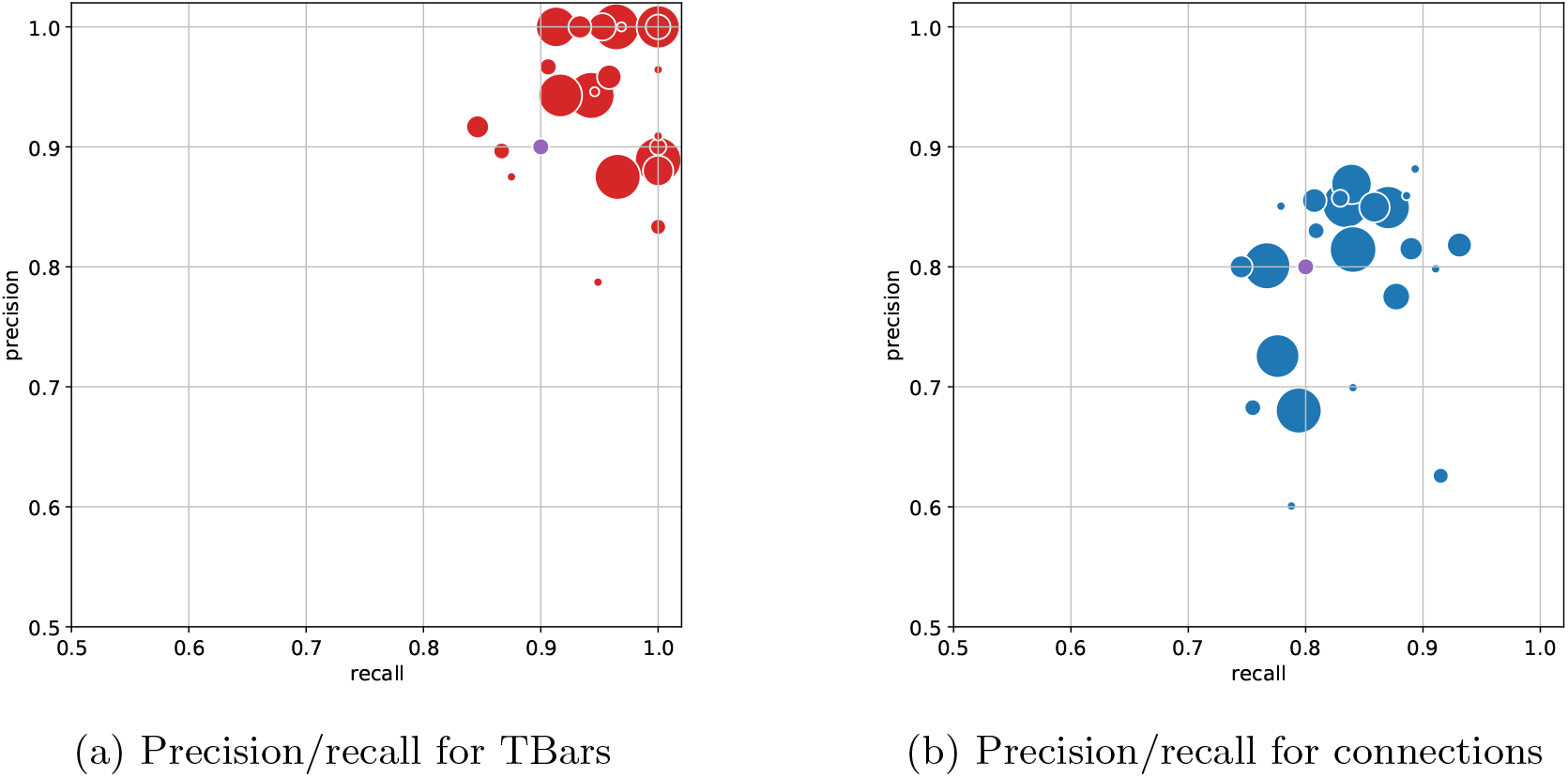
Precision/recall plots for synapse identification

### 2.7 Anatomical region definition

The synaptic neuropils were defined using synapse point clouds (as this was performed before segmentation). The region boundaries including the fiber bundles were hand-drawn based on the grayscale data. The standard nomenclature system for the VNC [Court et al., 2020] was used to identify the neuropils and fiber bundles, although some structures were not identifiable from the point cloud data.

### 2.8 Proofreading

Proofreading was largely performed as in the hemibrain[Scheffer et al., 2020]. The main difference was that in this sample, we assumed bilateral symmetry and therefore expected every neuron to have a symmetric partner. This enabled checks and error correction not possible in much of the hemibrain sample.

The primary completion metric used is called the completeness, which is the percentage of all synapses in a region where both the pre- and post-synaptic partners belong to identified neurons. The completion percentage per neuropil is shown in Table 1.

### 2.9 Light-EM correspondence

To facilitate comparisons between the EM and light microscopic data, we transformed the coordinates of the EM volume into the standard reference frame used for light microscopy. This was performed using image registration to compute a spatial transformation that warped the synapse cloud of the VNC onto the JRC2018M *Drosophila* male ventral nerve cord template [Bogovic et al., 2020].

The approach is similar to that described in [Scheffer et al., 2020]. Synapse predictions are rendered into an image volume at a resolution close to the JRC2018M VNC template (400nm isotropic), manually reoriented, and blurred with a 4×4×4 Gaussian kernel. Next, we ran the automatic image registration library elastix[Klein et al., 2010] in two steps, using the VNC template as a fixed image and the rendered synapse cloud as the moving image. The first step finds an affine transformation, and the second step is initialized with that affine and finds a nonlinear (B-spline) transformation. The result was further fine-tuned manually using BigWarp [Bogovic et al., 2016]. The transformed synapses from elastix, the raw EM, and the JRC2018M VNC template were jointly visualized, and 213 landmarks placed to define a thin plate spline transformation. The composition of the transformations from these steps is the spatial transformation from the JRC2018M VNC template to the VNC EM space. We also estimated the inverse of the total transformation using code found in https://github.com/saalfeldlab/template-building.

### 2.10 Neurotransmitter prediction

Computer vision prediction of neurotransmitter for each presynaptic location was performed using the procedures and software from [Eckstein et al., 2023] for the three most common neurotransmitters - acetylcholine, GABA, and glutamate. Ground truth neurotransmitter labels for 187 neurons (67 acetylcholine, 55 GABA, 65 glutamate) were annotated based on morphological identification with light level data. Neurons were partitioned into disjoint training and validation sets of 80% and 20%, respectively, optimizing for similar class frequency in each partition. Models included an additional class to indicate non-synaptic or unrecognized structures, trained by sampling random locations in the bounding box of all synaptic locations. Predictions from the validation-selected model were aggregated for each neuron as the mean of probabilities of all presynaptic sites. The maximum likelihood predicted probability for each neuron was annotated as the predicted neurotransmitter.

In NeuPrint, the computed probability of each neuro transmitter type is attached to the TBar component of each synapse. For a synapse *s*, these can be found as *s*.*ntAcetylcholineProb, s*.*ntGabaProb*, and *s*.*ntGlutamateProb*, and *s*.*ntUnknownProb* yields the probability that *s* matches none of those three. The value ranges from 0 (impossible) to 1.0 (certain). The values sum to 1 and should be interpreted as relative probabilities, as other transmitters are possible.

### 2.11 Hemilineage annotation

Segmentally repeating arrays of neuroblasts give rise to embryonic (primary) and postembryonic (secondary) neurons during development. Secondary neurons from each neuroblast are organized into one or two hemilineages previously described at light level, each with conserved neurotransmitter expression and similar morphology. These hemilineages therefore consist of developmentally related groups of neurons, which typically have similar morphology and functional properties. Hemi-lineages provide a natural high-level grouping for all of the neurons with cell bodies in the ventral nerve cord; we used this grouping extensively in all of the subsequent annotation and quality control procedures, including soma side and neuromere assignment, left-right matching, and cell typing. A full description of the procedures and outcomes is provided in Marin et al.

### 2.12 Soma side and soma neuromere annotation

Somas (cell bodies) were manually annotated early in the project, and the coordinates of their centroids were automatically assigned as a soma position value. Beginning with neurons that pass through the neck connective, neurons were clustered and then soma side and soma neuromere (or tract side for descending neurons) were automatically annotated by cluster, except that those near the midline were manually reviewed and annotated. Soma neuromeres in the abdomen were manually annotated by an expert. Both soma side and neuromere assignments were later refined based on hemilineage of origin and identification of groups and serial sets.

### 2.13 Group and serial set annotation

Several approaches were used to match similar neurons, including across the midline in the same neuromere, and assign them to groups. In addition to left-right groups, serial sets of homologous neurons were also identified by comparing morphology and connectivity across neuromeres, as described in Marin et al., and contributed to cell typing across VNC neuromeres.

Descending neurons and ascending neurons were matched by mirroring one side of the neck connective onto the other and performing an NBLAST clustering followed by manual review. Motor neurons were manually proofread and annotated with their exit nerve, then assigned to groups and serial sets by morphology and connectivity and matched to light level instances where possible. Full details of descending neuron and motor neuron matching and identification can be found in Cheong, Eichler, Stürner, et al.

The remaining neurons of VNC origin were assigned to groups using one of two approaches. The first approach was that used in the hemibrain, using NBLAST and CBLAST. In the second approach, a novel tool called NBLAST-DASH was developed to facilitate identification and co-visualization of neurons with similar morphology. NBLAST scores were used to cluster neurons sorted by hemilineage, soma side, and soma neuromere, generating a tanglegram that suggested likely matches across the midline. These candidate matches were visually compared by proofreaders and used to assign neurons to groups. The group assignments were later refined following proofreading and connectivity comparisons.

### 2.14 Cell typing and nomenclature

Assigning cell types to each reconstructed cell was a complex operation that interacted strongly with the grouping process described above as well as helping to ensure that neurons were reconstructed to the highest quality possible. Typing is more thoroughly described in Marin et al. This section provides a summary of that process.

Reconstructed neurons can be divided into four large categories: (1) Descending neurons with their somas in the brain and axons in the VNC; (2) Sensory neurons with their somas in the periphery; (3) Motor neurons with their somas in the VNC cortex and axons that exit the nerves to target muscles; (4) Other neurons with somas in the VNC cortex, comprising intrinsic neurons, ascending neurons, and efferent neurons. All neurons were initially annotated with a coarse class based on these distinctions and this was crucial to many analyses.

All neurons additionally received a finer scale *systematic type* name based only on our analyses of their morphology and connectivity within the MANC dataset. They also received a *type* name which is used in our companion manuscripts. In the majority of cases the *type* and *systematic type* are the same. However, for a minority of MANC neurons that have been clearly described in the prior literature and for which matches to suitable light level data could be made, the *type* name was drawn from existing systematic light level nomenclature. For certain well-known neurons for which multiple names exist, additional annotations were recorded in the *synonyms* field. Each neuron is also associated with an instance reflecting the nerve entry side for descending neurons and sensory neurons or soma side and neuromere for all other neurons, as well as a unique *bodyid* identifier.

In defining types, we used 4 slightly different formats for the coarse categories noted above to represent the most useful features concisely while maintaining consistency within each category. Complete details for identification and typing of descending neurons and motor neurons (total n=2065) are described in Cheong, Eichler, Stürner et al. Sensory neurons and intrinsic neurons of the VNC other than motor neurons (total n=21683) are described in detail in Marin et al. Intrinsic neurons were clustered into systematic types broader than left-right groups and serial homologue sets by a hierarchical clustering based on cosine connectivity similarity. Similarity was computed for connectivity symmetrized across left-right groups and across serial homologue sets, while parameters for final clusters were selected based on consistency between left and right lateral populations considered independently. Hemilineage and serial set annotations were used to constrain clusterings and linkage between clusters, respectively. Complete details and analysis for this process and the resulting systematic types are provided in Marin et al.

### 2.15 Concluding Remarks and Future Research

The MANC is a milestone in connectomics. It captures more synaptic connectivity than any connectome yet released. It is the first complete nerve cord connectome and the first connectome of a bilaterally complete region of the central nervous system of an adult animal. For these reasons the MANC connectome will be of significant interest to computational neuroscientists, graph theorists and others using quantitative methods to investigate the nervous system. For neurobiologists the MANC dataset has been richly annotated with cell type information at different granularity and also benefits from neurotransmitter predictions across the dataset. This will immediately allow existing hypotheses to be refuted or supported and many new hypotheses to be generated.

This connectome will also play a key role in study of stereotypy of neuronal wiring in *Drosophila*. The sample was one of a set of flies, with identical genetic backgrounds and experience, that were raised together and eventually simultaneously prepared, fixed, and preserved. Another of these samples is used in the upcoming male central nervous system, which will also include the VNC of that sample. This will provide for the first time two large, densely proofread, examples from two identical specimens, enabling study of the inherent variation between animals.

We carry out a preliminary analysis of important features of the biological organization of the VNC in our two companion papers. In Marin et al, for the first time, we provide a comprehensive link between the developmental origins of VNC neurons (from different neural stem cells, called neuroblasts) and their circuit properties defined by the connectome. We also reveal at single neuron precision the extensive serial homology between different segments of the nerve cord, especially between the circuits controlling the legs. In Cheong, Eichler, Stürner et al we provide a more detailed characterisation of the organization of the circuits between approximately 1300 descending neurons and 700 motor neurons. We identify a serially repeated standard leg connectome and the intersegmental interneurons that likely coordinate movement across the legs. We describe key circuit features of flight control and compare and contrast the pre-motor organization of leg and wing circuits. We also provide initial investigations of circuits associated with turning and reversing while walking, take-off, flight and song. These provide a platform for detailed further studies of these and many other behaviors which we expect to follow rapidly from many different laboratories. Crucially, these initial investigations ensure that the base connectome, annotations and analysis tools are validated and ready for use by the wider research community.

## 3 Acknowledgements

This work was supported by the Howard Hughes Medical Institute and the Wellcome Trust (220343/Z/20/Z and 221300/Z/20/Z to GSXEJ with GMR, GMC, Scott Waddell and Matthias Landgraf). Portions of this work were performed by the FlyEM Project Team at Janelia Research Campus, in addition to the named authors. We thank Barry Dickson and his lab for providing ground truth data used for neurotransmitter prediction.

## 4 Author Contributions

KJH and ZL developed sample preparation and prepared the sample; HFH and CSX developed the imaging hardware; CSX imaged the sample; KH, DaK, MN, CO, SP, SS, LS, and ET aligned the images; SB, MJ, CO, and ShT created the segmentation; GH, CO, and PR found the synapses; JB performed the LM-EM mapping; ECM, CO, and KS determined the neuropil and nerve regions; AC, JF, and GSXEJ predicted neurotransmitters; GB, DB, SB, BC, HC, MiC, OD, CD, K, SF, SFM, AF, MG, GPH, JH, CK, JK, SL, FL, AL, ChM, EM, ECM, CaM, BM, EN, OO, NO, CO, TP, ElP, EmP, CR, PR, SR, TR, AKS, ALS, AyS, CS, NS, AlS, SaT, ShT, IT, IFMT, JW, and TY did the proofreading; GB, SB, PB, GMC, HC, MC, CD, KE, SF, MG, JH, AJ, GSXEJ, CK, DoK, FL, ECM, BM, EN, CO, SMP, PR, SS, DS, TS, ShT, IFMT, ET, and LU did quality control; GB, PB, GMC, AC, HC, MC, CD, KE, MG, GSXEJ, DoK, FL, ECM, BM, DS, TS, ShT, and IFMT performed cell typing; SB, JB, JC, KH, GH, PH, MJ, GSXEJ, DaK, WTK, DoK, BM, MN, SMP, SP, SS, PS, ET, LU, and TZ contributed software; GMC, MJ, GSXEJ, WK, ECM, GMR, and LS wrote the paper; SB, GMC, MC, KE, RG, HFH, GSXEJ, CK, FL, ECM, EN, SMP, SP, PR, and SS coordinated internal efforts; ECM coordinated annotation; GSXEJ managed the Cambridge effort; SB managed the overall effort.

